# Myo1e modulates the recruitment of B cells to inguinal lymph nodes

**DOI:** 10.1101/668608

**Authors:** Daniel Alberto Girón-Pérez, Eduardo Vadillo, Michael Schnoor, Leopoldo Santos-Argumedo

## Abstract

The recruitment of leukocyte to high endothelium venules and their migration to the lymph nodes are critical steps to initiate an immune response. Cell migration is regulated by the actin cytoskeleton where myosins have a very import role. Myo1e is a long tail class I myosin highly expressed in B cells that not have been studied in the context of cell migration. By using an *in vivo* model, through the use of intravital microscopy, we demonstrated the relevance of Myo1e in the adhesion and the migration of B cells in high endothelial venules. These observations were confirmed by *in vitro* experiments. We also registered a reduction in the expression of integrins and F-actin in the protrusion of B lymphocytes membrane. Deficiencies in vesicular trafficking can explain the decrease of integrins on the surface. Interestingly, Myo1e is associated with focal adhesion kinase (FAK). The lack of Myo1e affected the phosphorylation of FAK and AKT, and the activity of RAC-1, disturbing the FAK/PI3K/RAC-1 signaling pathway. Together, our results indicate critical participation of Myo1e in the mechanism of B cell migration.

**Summary statement:** Myo1e participate in the adhesion and migration in the high endothelial venules by regulation of integrins and the PI3K/FAK/RAC-1 signaling pathway.

## Introduction

The secondary lymphoid organs have a critical role in immunity. Their distribution in the body allows the recruitment of immune cells for encountering antigens (Okada and Cyster, 2006, Pereira et al., 2010, Mesin et al., 2016). The lymphocytes adhere to high endothelial venules (HEV) for crossing to lymph nodes to look for their antigen and to mount an immune response. The adhesion and migration are mediated by highly controlled mechanisms regulated by integrins, adhesins, chemokines and the actin cytoskeleton (Anderson and Anderson, 1976, Mionnet et al., 2011, Girard et al., 2012). The dysregulation of these molecules can cause a reduction in migration and recruitment of lymphocytes affecting the immune response. Therefore, it is necessary to analyze the elements in detail to better understand the mechanism of migration.

Cell migration consists of various steps highly regulated by signaling molecules (i.e., GTPases, kinases or motor proteins) (Mayor and Etienne-Manneville, 2016, Vicente-Manzanares et al., 2005, De Pascalis and Etienne-Manneville, 2017, Mitchison and Cramer, 1996) that control morphological changes needed for the movements of the cells (Mitchison and Cramer, 1996). These changes, modulated by alterations in the cytoskeleton, control the extensions of their plasma membrane (Santos-Argumedo et al., 1997, Maravillas-Montero et al., 2011). Myosins are motor proteins included in 18 families (Thompson and Langford, 2002) and expressed by different tissues and organisms (Sellers, 2000). Class I myosins are single head molecules that can bind to the actin filaments and the plasma membrane (Osherov and May, 2000). The functions of class I myosin are associated with the regulation of motility and adhesion. Myo1e is highly expressed by macrophages, dendritic cells, and B cells (Santos-Argumedo et al., 2013, Wenzel et al., 2015). In macrophage and dendritic cells, Myo1e have a role in antigen presentation due to the association of Myo1e with ARF7EP (Paul et al., 2011). The absence of Myo1e affects the transport of MHC-II to the plasma membrane Additionally, the lack of Myo1e in activated macrophages reduce their cellular spreading (Tanimura et al., 2016). In the case of Myo1f, studies using intravital microscopy have shown the relevance of this protein in the extravasation, migration, and deformation of the nucleus of neutrophils (Salvermoser et al., 2018). While, in infections with *Listeria monocytogenes*, the motility of neutrophils is reduced (Kim et al., 2006). Given, the capacity of long tail class I myosin in regulating the migration and adhesion, this study focused on the evaluation of Myo1e during B cells migration. Here we report that the long tail Myo1e participates in the adhesion and slow rolling of B cells in HEV of the inguinal lymph node. By *in vitro* assays, we demonstrated that the loss of Myo1e causes a reduction in the expression of integrins in the membrane and is associated with the PI3K/FAK/RAC-1 signaling pathway. Thus, these results indicate the critical participation of the Myo1e in the process of migration and its possible functions in the regulation of adhesion and extravasation.

## Results

### In the absence of Myo1e, there is inefficient recruitment of B cells to the inguinal lymph node

The recruitment of leukocytes to the lymph nodes is a critical step for mounting an immune response; this process involves the rolling and adhesion of leukocytes to the venules of the high endothelium (Kansas et al., 1993). We investigated whether Myo1e, expressed by B cells (Santos-Argumedo et al., 2013, Maravillas-Montero et al., 2011) affects the adhesion and the motility of these cells to the venules of the high endothelium. Therefore, the migration of activated B cells from control mice (Myo1e^+/+^) was compared with Myo1e-deficient mice (Myo1e^−/−^). Hoescht 33342-labeled B cells were injected into a host wild type mouse. The adhesion, rolling and migration was evaluated by intravital microscopy in the inguinal lymph node using CXCL12 as a chemoattractant.

First of all, the venules of the lymphatic node of the host mouse were evaluated, where the parameters proposed by Von Adrian UH in 1996 were analyzed (Von Andrian, 1996), Those parameters include the numbering of the branches of the inguinal lymph node from I to IV (Fig.S1), as well as the diameter and blood flow of the different venules (Fig. S2A-B). Subsequently, the migration of B cell in the absence of any additional chemokine (vehicle), was registered for both the B cells obtained from control mice and Myo1e^−/−^ mice. In both situations, the adhesion and migration were negligible (Video Movie 1-2, (Duration 00.29 seconds). In sharp contrast, when the chemoattractant CXCL12 was used, lower adhesion and reduced migration were observed for B cells originated from Myo1e-deficient mice (Myo1e^−/−^) when compared with control mice (Myo1e^+/+^) (Video Movie 3-4) (Duration 00:29 seconds).

### Myo1e is essential for the recruitment and adhesion of B cells to the inguinal lymph node

Derived from the previous observations, the recruitment of B cells, in the absence of Myo1e, was investigated in further detail. As can be seen in Fig. 1A-B, there is a reduction in the recruitment of B cells from Myo1e^−/−^ mice, regardless of time. Additionally, there is an increase in cell flow in venules I and II in Myo1e^−/−^ B cells compared with the control mice (Myo1e^+/+^) (Fig. 1C-D). In correspondence with the previous results, we registered reduced adherence of Myo1e^−/−^ B cells to the III and IV venules. These venules contain high endothelial cells (Fig. 1E). These results suggest that Myo1e modulates the migration of B cells into the inguinal lymph node.

**Fig 1.**
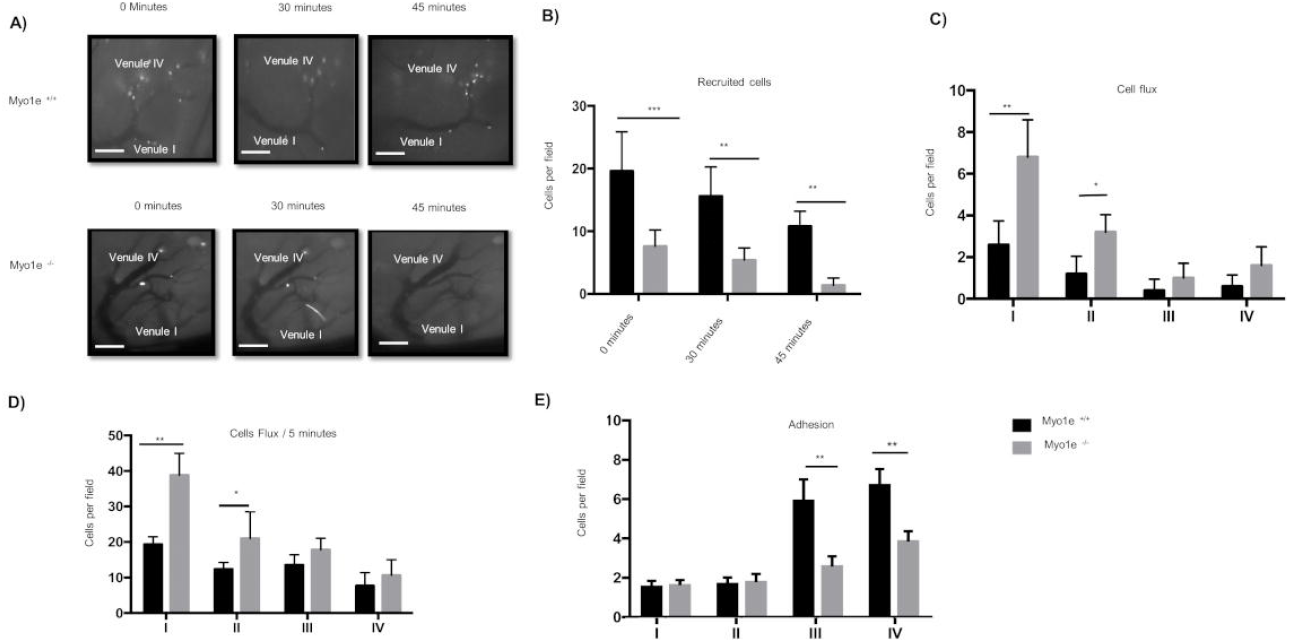
Myo1e is required for recruitment and adhesion of B cells to the inguinal lymph node. A) Representative images of intravital microscopy of activated B cells (stained with Hoescht 33342) from Myo1e^+/+^ and Myo1e^−/−^ mice in the venules of an inguinal lymph node of a host Myo1e^+/+^ mice. Images were registered (40x objective) at different time points (0, 30 and 45 minutes) in the venules of an inguinal lymph node that was previously injected (1 h) with CXCL12 (25 ng/ml) or the vehicle (PBS). The venules were identified as IV to I. Scale bar: 25 μm; n=5. B) Quantification of recruited B cells C) Measurements of B cell flux. D) Measurements of B cell flux at 5 minutes. E) Numbers of adherent B cells. The quantifications were performed in the different venules (IV to I), n=5. Data are represented as mean ± SEM. *** p<0.001, **p<0.01 *p<0.05.

### The deficiency of Myo1e affects the speed and the slow rolling of B lymphocytes

To follow with the characterization of the motility of B lymphocytes, we evaluated the time it takes to B cells of travel from venule IV to I (Fig. 2A). We registered an increase in the rolling of the B cells from Myo1e^−/−^ mice traveling in venules IV and III, compared with B cells from wild type mice (Fig. 2B). In contrast, the analysis of “slow rolling” (frequency of leukocytes with a rolling velocity less than 5 μm/sec), (Weninger et al., 2000), showed a reduction in this parameter by B cells from Myo1e^−/−^ mice (Fig. 2C). Both results reflect an increase in the speed of B cells from Myo1e^−/−^ mice compared with Myo1e^+/+^ mice (Fig. 2D). These results agree with the reduction in cellular transmigration (Fig S3) and indicate that Myo1e participates in the adherence and rolling of B lymphocytes to HEV.

**Fig 2.**
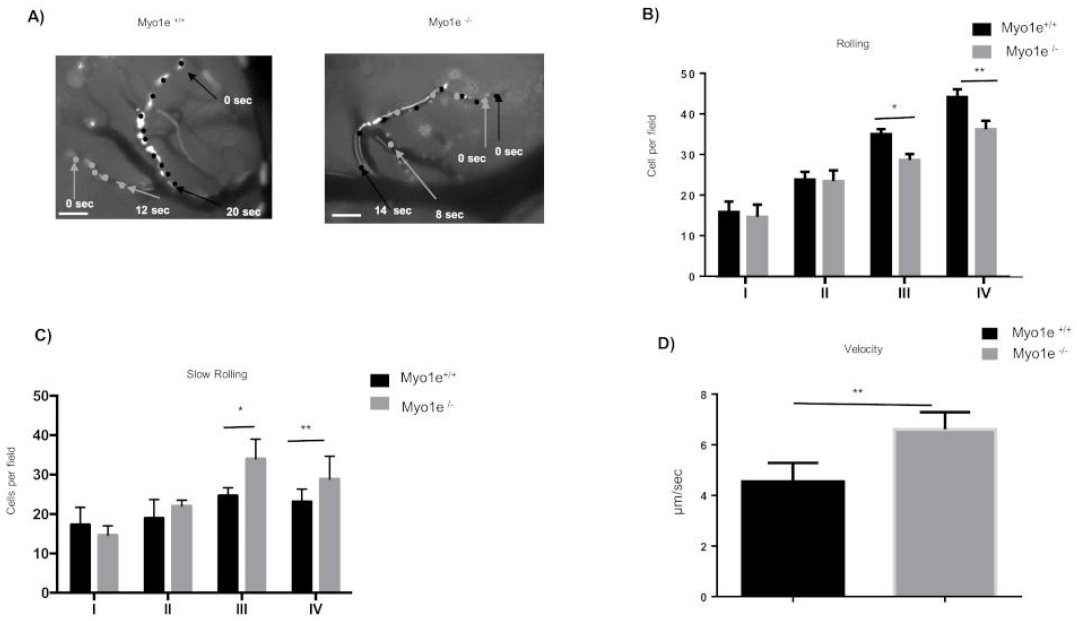
The lack of Myo1e causes a reduction in the slow rolling and the velocity of activated B lymphocytes. A) Representative images of the migration of activated B cells (stained with Hoescht 33342) from Myo1e^+/+^ and Myo1e^−/−^ mice in the venules of an inguinal lymph node of a host Myo1e^+/+^ mice. The inguinal lymph node was previously injected (1 h) with CXCL12 or vehicle (PBS). The arrows indicate the start of the route of B cells from venule IV to I (40x objective). Scale bars 25 μm; n=5. B) Quantification of the numbers of activated Myo1e^+/+^ and Myo1e^−/−^ B cells performing rolling in the different venules (IV to I) of an inguinal lymph node of a host Myo1e^+/+^ mice. n=5. Data are represented as mean ± SEM.*** p<0.001, **p<0.01 *p<0.05. C) Quantification of numbers of activated Myo1e^+/+^ and Myo1e^−/−^ B cells performing slow rolling in the different venules (IV to I) of inguinal lymph node of host Myo1e^+/+^ mice n=5. Data are represented as mean ± SEM.*** p<0.001, **p<0.01 *p<0.05. D) Measurements of the velocity of displacement of activated B cells from of Myo1e^+/+^ and Myo1e^−/−^ from venule IV to I in the inguinal lymph node; n=5. Data are represented as mean ± SEM. *** p<0.001, **p<0.01 *p<0.05.

### The loss of Myo1e in B cells affects their CXCL12-dependent homing to the inguinal lymph node

To corroborate our findings, homing assays were performed in which the right lymph node was inoculated with CXCL12, while the left lymph node was injected with the vehicle (Fig. S4). Subsequently, CFSE-labeled activated B cells from Myo1e^+/+^ and Myo1e^−/−^ mice were injected in different cell proportions into a host wild type mouse. We found a reduction in the recruitment of Myo1e deficient B cells in the right, but not in the left lymph node (Fig. S4B). To further corroborate these results, images were taken by intravital microscopy where, compared with the right inguinal lymph node, and left lymph node (Supplementary Fig. 4C-D). In contrast, we found more Myo1e^−/−^ B cells recirculating in the blood and the spleen, indicating that those cells were not recruited into the lymph node (Fig S5). These findings demonstrate that Myo1e is critical to modulate the process of B cell migration.

### Myo1e modulate the chemotaxis of B lymphocytes

To analyze how the absence of Myo1e affect chemotaxis, the migration of resting and activated B lymphocytes was evaluated in a Zigmond chamber. Myo1e-deficient activated B cells showed reduced trajectories in comparison with activated B cells from wild type control mice (Fig. 3A). This deficiency was also reflected in the direction ratio (angles that makes a straight line when changing directions) (Fig. 3B), Euclidian distance (straight line distance between two points), accumulated distance (total distance traveled between two points) (Fig. 3C); and, the velocity (Fig. 3D). In the experiments, using resting B cells, we did not find any difference (Fig. S6). Because the differences were only found using activated B lymphocytes, the following experiments were performed using these cells. These results indicate the critical relevance of Myo1e in the migration.

**Fig. 3.**
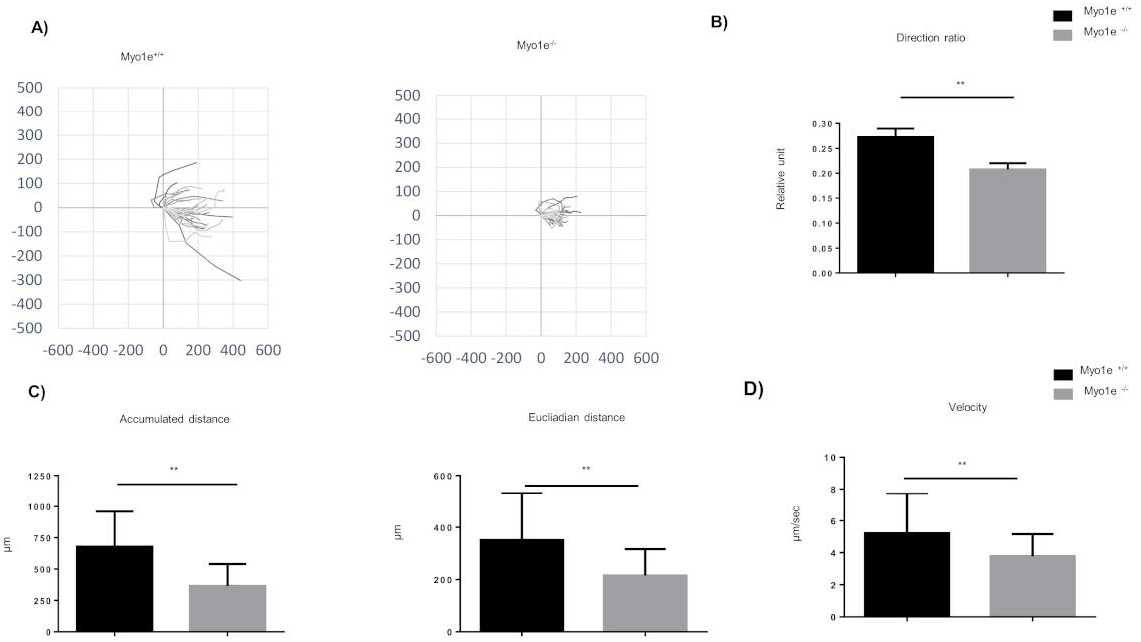
The absence of Myo1e affects the distance and the 2D motility in response to CXCL12. A) Activated B cells from of Myo1e^+/+^ and Myo1e^−/−^ mice were deposited in the Zigmond chamber under a CXCL12 gradient and registered for 1 hour. Tracks of individual trajectories are presented in the plots; n=5. B) Measurement of the direction ratio. C) Quantification of the accumulated and the Euclidian distances. D) Measurements of the velocity. The experiments were performed with resting or activated Myo1e^+/+^ and Myo1e^−/−^ B cells under a CXCL12 gradient; n=5. Data are represented as mean ± SEM. *** p<0.001, **p<0.01 *p<0.05.

### Myo1e regulates the expression of integrins and adhesion molecules, affecting cell adhesion

Integrins and adhesion molecules strongly modulate cell migration in different cells and tissues (Walling and Kim, 2018, Senbanjo and Chellaiah, 2017, Chuluyan and Issekutz, 1993, Manevich et al., 2007, Smith et al., 2003, Gerberick et al., 1997). These molecules allow the adherence of the cells to the extracellular matrix, which serves as a support for the elongation of the membrane to generate the force needed for motility (Francois et al., 2016, Sales et al., 2019, Doyle et al., 2015). Due we found defects in the motility of Myo1e-deficient B cells, we measured how were the relative amount of LFA-1, CD44, and VLA-4 in activated B cells from Myo1e^−/−^ B cells compared with the control wild type mice. The analysis of the mean fluorescence intensity (MFI) in activated B cells from Myo1e^−/−^ mice showed a reduced amount of LFA-1, CD44, and VLA-4 on the membrane of these mice compared with B cells from Myo1e^+/+^ mice (Fig. S7A). In the experiments, using resting B cells, we did not find any difference (Fig. S8). Also, when the full expression of these adhesion molecules was analyzed, no significant differences were found; other proteins, such as CXCR4, TLR-4 and CD62-L did not show these differences (Fig. S9). The decrease in LFA-1, CD44, and VLA-4. In adhesion assays using a monolayer of b.End3 cells the activated B cells Myo1e^−/−^ mice, have a reduced capacity of adhesion in comparison with the Myo1e^+/+^. (Fig. S7B-C)). These data suggest that Myo1e modulates cell migration through controlling the expression of LFA-1, CD44, and VLA-4 on the surface.

### Cell transmigration and membrane prolongation requires the presence of Myo1e

We next evaluated adhesion on specific substrates (fibronectin, hyaluronate acid and poly-L-Lysine as control). Myo1e-deficient activated B lymphocytes showed a significative reduced adherence to fibronectin and hyaluronic acid compared with B cells from Myo1e^+/+^ mice (Figure. 4A). The adhesion to a non-specific substrate like poly-L-Lysine did not show differences. In the experiments, using resting B cells, we did not find any difference (Fig. S10A). To extend this observation, we analyzed cellular transmigration through monolayers of b.End3 cells (brain endothelial cell line from SV129 mice) in a trans-well chamber. We found a reduction in the transmigration of activated B cells when Myo1e was absent, but not in resting B cells (Fig. 4B and Fig. S10B). Interestingly, when measuring the membrane extensions in activated B cells, Myo1e^−/−^ B cells exhibited reduced membrane prolongations (Fig. 4C-D). These observations agree with a defective migration of B lymphocytes from Myo1e^−/−^ mice due to their reduced adhesion to the substrate.

**Fig. 4.**
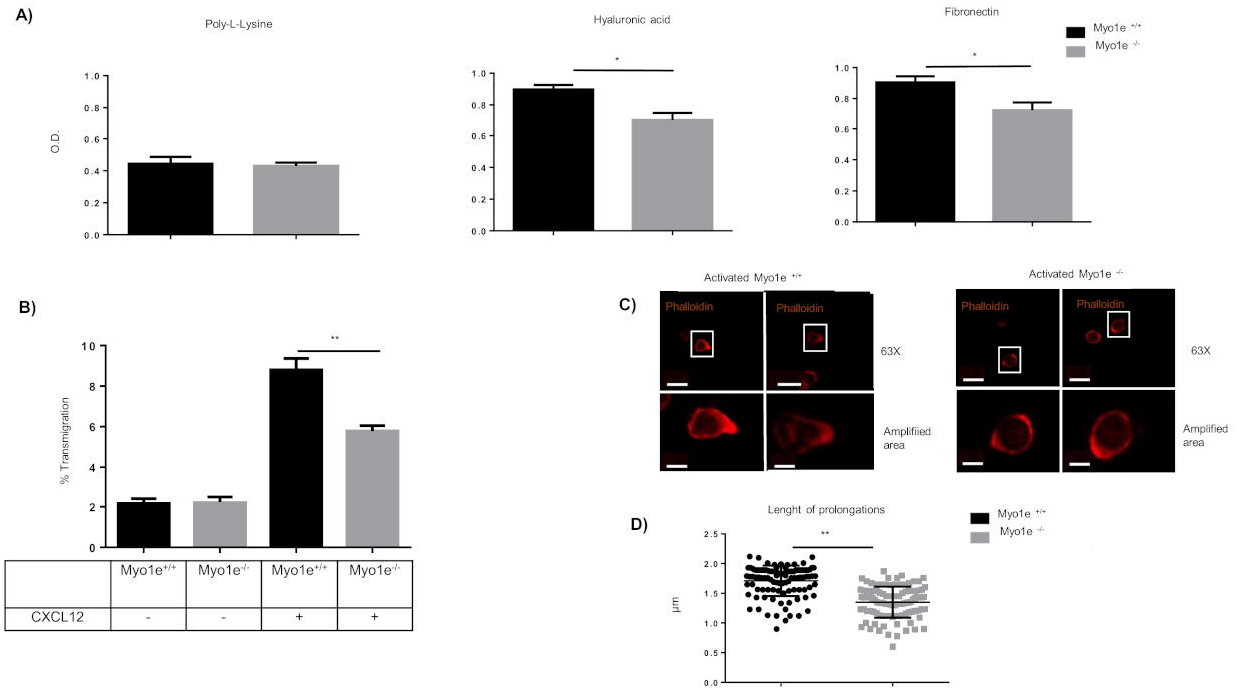
The deficiency of Myo1e affects the transmigration and the length of protrusions of the membrane of activated B cells. A) One hundred thousand activated B cells from Myo1e^+/+^ and Myo1e^−/−^ mice were placed into each well in a 96 wells plate. Previously, the plate was coated with hyaluronic acid, fibronectin or poly-L-lysine for two hours; then, the wells were washed, and the cells adhered to the wells were stained with crystal violet. Finally, the cells were lysed, and the absorbance of the dye was determined at 590 nm; n=3. Data are represented as mean ± SEM. *** p<0.001, **p<0.01 *p<0.05. B) Activated B cells from of Myo1e^+/+^ and Myo1e^−/−^ mice were seeded in a trans-well chamber under a gradient of CXCL12 or only medium for four hours. Previously, the chambers were seeded with bEnd.3 cells until they formed a monolayer (usually two days). Migrating B cells were recovered from the bottom chamber and quantified by flow cytometry. Percentages of transmigration are presented in the graph; n=3. Data are represented as mean ± SEM. *** p<0.001, **p<0.01 *p<0.05. C) Representative images (63x objective) of activated Myo1e^+/+^ and Myo1e^−/−^ B cells under a gradient of CXCL12. Scale bars 5 μm; n=3. D) Measurement of the length of protrusions of activated Myo1e^+/+^ and Myo1e^−/−^ B cells. Data are represented as mean ± SEM. *** p<0.001, **p<0.01 *p<0.05.

### The localization of integrins in the membrane protrusions is affected by the absence of Myo1e and FAK is physically and functionally associated with it

To determine how Myo1e is involved in the signaling mediated by integrins, the pixel intensity of LFA-1 in the protrusion was measured. We found a lower signal intensity in the protrusions of B lymphocytes from Myo1e^−/−^ mice, (Fig. 5A); in contrast, the expression of CXCR4 was not reduced (Fig S11A-B).

**Fig. 5.**
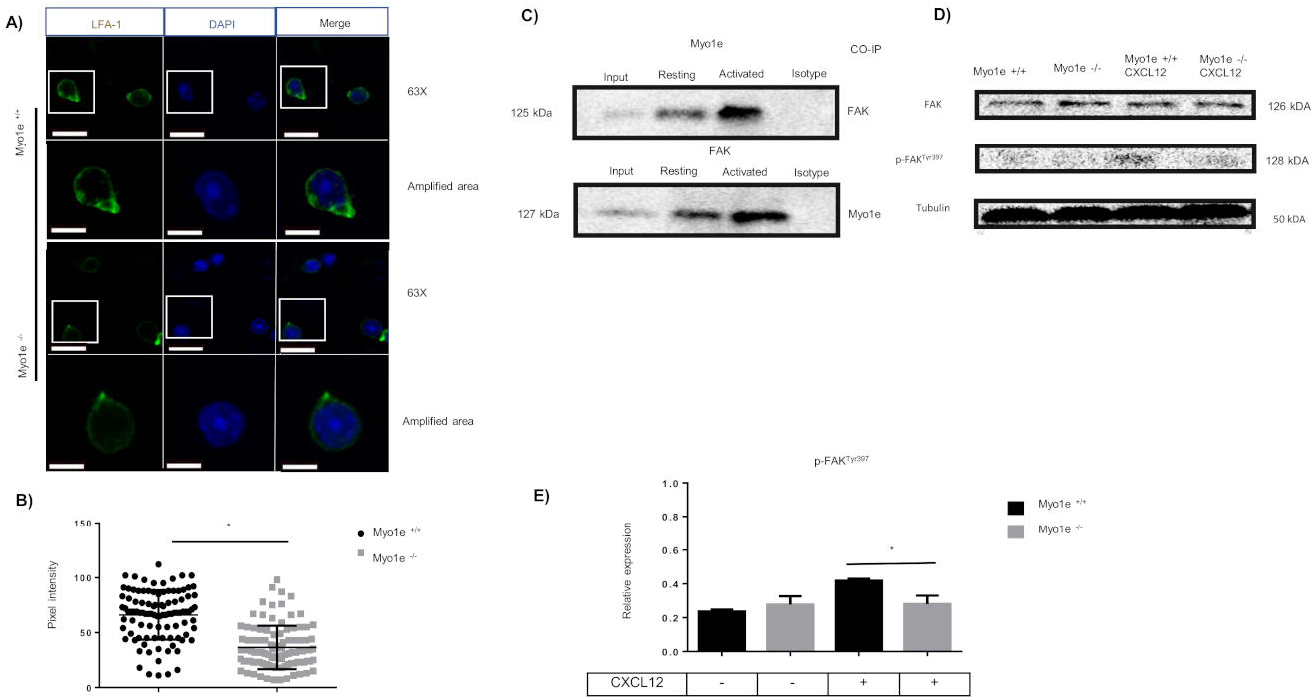
Myo1e interacts with FAK and the lack of Myo1e causes a reduction in the localization of integrins. A) Representatives images (63x objective) of activated B cells from of Myo1e^+/+^ and Myo1e^−/−^ mice, under a CXCL12 gradient. The cells were stained with anti-LFA-1 (green) and DAPI (Blue). Scale bars 5 μm. B) The intensity of pixels in the protrusion of membrane of activated B cells from of Myo1e^+/+^ and Myo1e^−/−^ mice; was measured n=3. Data are represented as mean ± SEM. *** p<0.001, **p<0.01 *p<0.05. C) Co-immunoprecipitation of Myo1e with focal adhesion kinase (FAK) in resting and activated B cells; n=3. D) Western blot of the Tyrosine 397 phosphorylation of FAK in activated B cells with or without stimulation with CXCL12; n=3. E) Densitometric analysis of tyrosine 397 phosphorylation of FAK; n=3. Data are represented as mean ± SEM. *** p<0.001, **p<0.01 *p<0.05

Because the focal adhesion kinase (FAK) is a protein that plays a critical role in integrin-mediated signal transduction, we looked if there was an association between FAK and Myo1e. We also searched for the phosphorylation at tyrosine 397 of FAK in B cells that were stimulated with CXCL12. The results showed that Myo1e could be co-immunoprecipitated with FAK. This association is stronger in activated Myo1e^+/+^ B cells (Fig 5B). Interestingly, FAK^Y397^ become phosphorylated in activated wild type B cells that were stimulated with CXCL12 (Fig. 5C-D). The phosphorylation is higher in wild type B cells compared with Myo1e^−/−^ B cells. These results showed a physical and functional association between FAK and Myo1e, suggesting that Myo1e is enclosed in the signaling pathway of integrins.

### Myo1e interacts with CARMIL affecting the polymerization of actin in the membrane protrusions

We evaluated the polymerization of actin in the protrusions of migrating B lymphocytes, finding a reduction of F-actin at the leading edge of the membrane in Myo1e^−/−^ B cell (Fig S12A-B). The reduction was specific to this site because we did not find differences in total actin (Fig S13).

CARMIL (capping protein, Arp2/3, and Myosin-I linker) is a family of proteins involved in the migration of the cells. CARMIL was analyzed through colocalization assays (Fig S12C-D), and by co-immunoprecipitation (Fig S12E). Both strategies showed that Myo1e and CARMIL are associated at the leading edge of migrating B lymphocytes.

These results indicate that Myo1e participates in the remodeling of filamentous actin at the leading edge of migrating B lymphocytes.

### Myo1e deficiency affects cellular spreading; besides, it affects the activity of RAC-1 and the phosphorylation of AKT

The Rho family of small GTPases are key regulators of the actin cytoskeleton and controlling the activity of numerous downstream effectors. Rac1, a member of the Rho family, together with AKT (a serine/threonine kinase) participate in actin reorganization required for the formation of protrusions during adhesion, spreading and motility of the cells. Thus, we evaluated the cellular spreading by measuring the curvature of the cell. If the ratio of semi-major ratio versus semi-minor axis (elliptical factor) > 2, this was indicative of a cell with polarized morphology. Most Myo1e^−/−^ B cells had an elliptical factor less than two compared with wild type B lymphocytes (Fig. 6A-B).

**Fig. 6.**
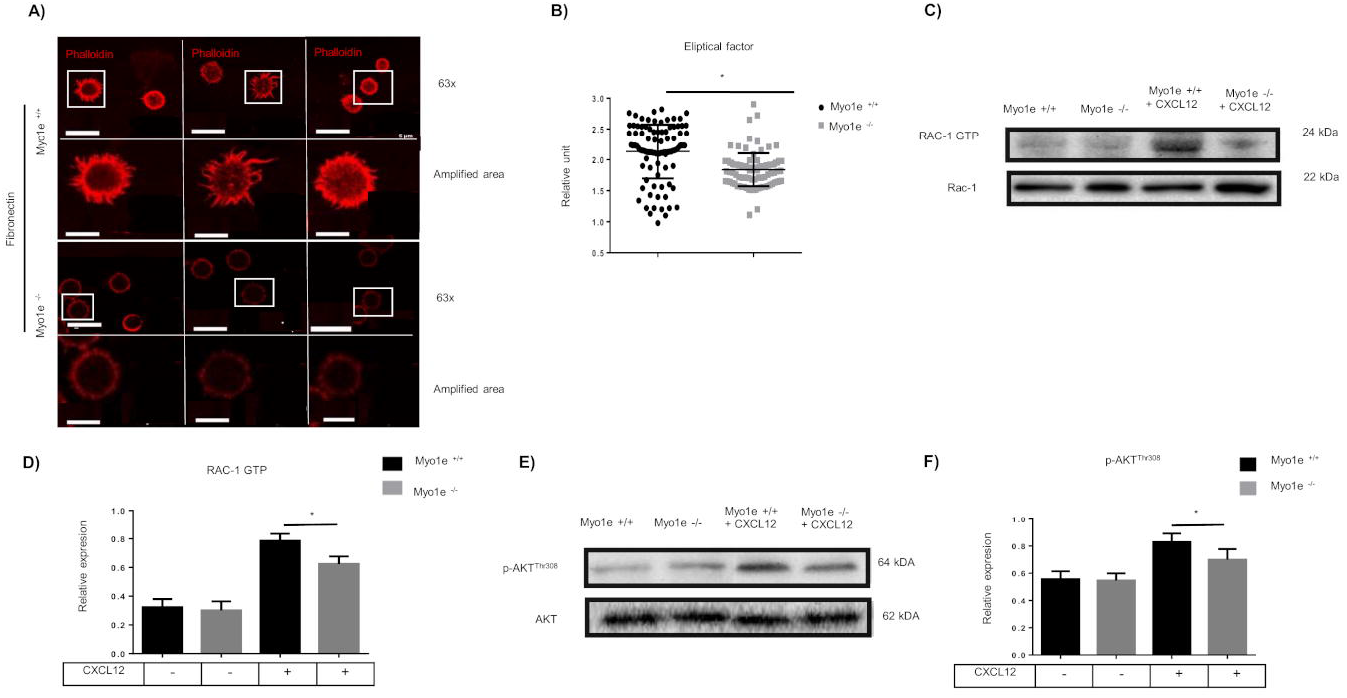
Myo1e is critically required for spreading and requires the activation of AKT/RAC-1 pathway. A) Representatives images (63x objective) of activated B cells from Myo1e^+/+^ and Myo1e^−/−^ mice. B cells were seeded to spread over fibronectin for 1 hour, and then, they were stained with TRITC-Phalloidin. Scale bars 5 μm; n=3. B) Quantification of the elliptical factor in Myo1e^+/+^ and Myo1e^−/−^ B cells; n=3. Data are represented as mean ± SEM. *** p<0.001, **p<0.01 *p<0.05. C) Western blot of the active form of RAC-1 in activated Myo1e^+/+^ and Myo1e^−/−^ B cells, with or without stimulation of CXCL12; n=3. D) Densitometric analysis of the activity of RAC-1; n=3. Data are represented as mean ± SEM. *** p<0.001, **p<0.01 *p<0.05. E) Western blot of Threonine 308 phosphorylation of AKT in activated B cells from of Myo1e^+/+^ and Myo1e^−/−^ mice, with or without stimulation of CXCL12; n=3. F) Densitometric analysis of Threonine 308 phosphorylation of AKT; n=3. Data are represented as mean ± SEM. *** p<0.001, **p<0.01 *p<0.05.

The evaluation of RAC-1 in Myo1e-deficient B cells demonstrated a decrease in their activity of this GTPase (Fig. 6C-D). Similarly, the phosphorylation of Threonine 308 of AKT was reduced in Myo1e-deficient B lymphocytes (Fig. 6E-F). These results show the relevance of Myo1e in the activity of actin-related proteins such as GTPase RAC-1 and AKT (Niba et al., 2013, Zhu et al., 2015, Henderson et al., 2015).

### Myo1e participates in controlling the migration of B lymphocytes through the signaling pathway of PI3K/AKT/RAC-1

The regulation of AKT and Rac-1 is dependent on the activation of PI3K (lipid kinase that phosphorylated lipids). Both enzymes are needed for elongation of the membrane driven by F-actin at the leading edge. By using a PI3K inhibitor LY294002, we found a decrease in the elongation and reduced F-actin at the leading edge. The results strongly resemble those seen with Myo1e-deficient B cells (Fig. 7A-C). These results correlate with the reduction of FAK^Y397^ and AKT^Thr308^ phosphorylation and RAC-1 activity (Fig. 7D and Figure S14). As a whole, these results strongly suggest that Myo1e is critical for cell migration and integrins activation. The FAK/AKT/RAC-1 signaling pathway requires the participation of Myo1e.

**Fig. 7.**
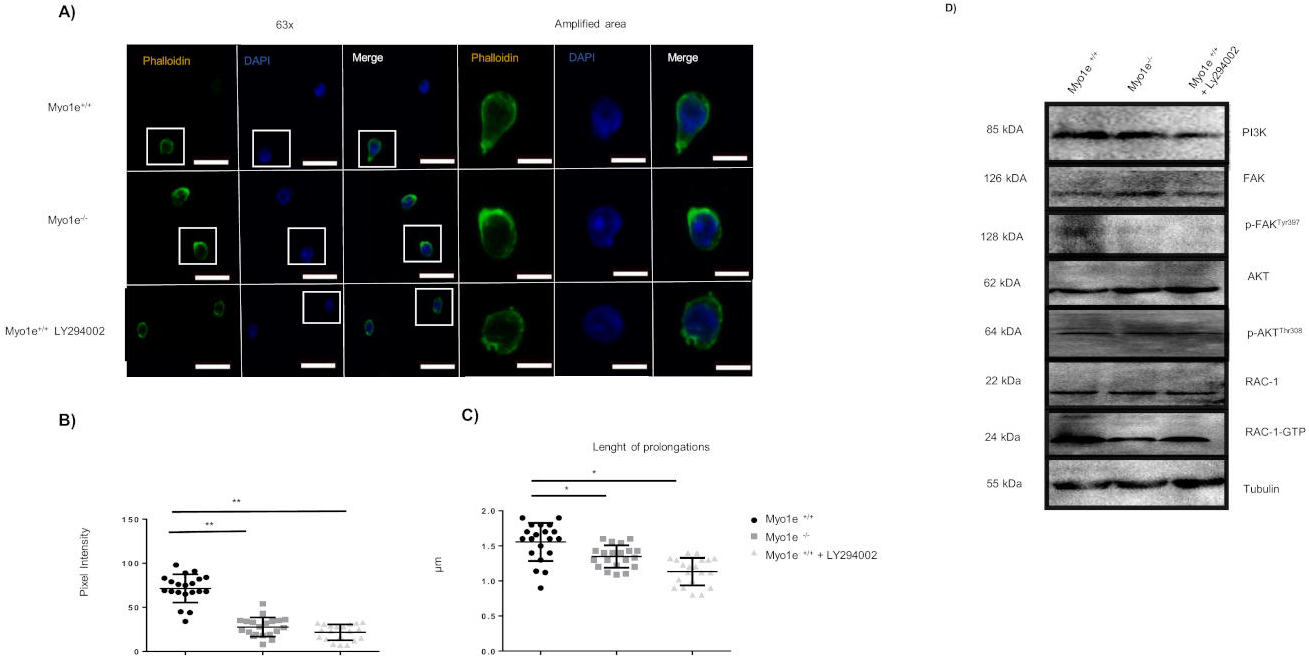
The inhibition of PI3K affects the protrusion of the membrane of activated B cells and requires the FAK/PI3K/AKT pathway. Representative images (63x objective) of activated B cells from Myo1e^+/+^ and Myo1e^−/−^ mice under a gradient of CXCL12 and treatment with LY294002. Scale bars 5 μm; n=3. B) Pixel intensity in the protrusion of the membrane of activated Myo1e^+/+^ and Myo1e^−/−^ B cells, under a gradient of CXCL12 and treatment with LY294002; n=3. Data are represented as mean ± SEM. *** p<0.001, **p<0.01 *p<0.05. C) Length of the protrusion of the membrane in activated Myo1e^+/+^ and Myo1e^−/−^ B cells under a gradient of CXCL12 and treatment with LY294002. All data shown are representative of three independent experiments performed. Data are represented as mean ± SEM. p<0.05. D) Western blot of PI3K and p-Akt (Thr308) and p-FAK (Tyr397) phosphorylation and activity of RAC-1 in activated Myo1e^+/+^ and Myo1e^−/−^ B cells under a gradient of CXCL12 and treatment with LY294002; n=3. Data are represented as mean ± SEM. *** p<0.001, **p<0.01 *p<0.05.

## Discussion

Class I myosins have been involved in the regulation of adhesion, motility and the recycling of receptors through the transport of vesicles and by the interaction with different cytoskeletal proteins (Piedra-Quintero et al., 2019, López-Ortega and Santos-Argumedo, 2017, Maravillas-Montero et al., 2011). However, few works are analyzing how class I myosins participate in the signaling mechanism of in leukocytes during cell migration (Salvermoser et al., 2018, López-Ortega et al., 2016). In the present study, through intravital microscopy, we demonstrated the relevance of the Myo1e in the motility of activated B cells. The loss of Myo1e causes a reduction in B cell recruitment to the inguinal lymph nodes. This reduction is accompanied by a decrease in slow rolling, as well as adhesion in high endothelial venules (HEVs). Similar results have been reported in Myo1f-deficient neutrophils, causes a reduction in spreading, transmigration and extravasation in cremasteric venules. This deficiency is due to an alteration in the morphology of the cell that prevents it from adhering correctly and allowing it to transmigrate through the tight junctions; however, the mechanism of action is not discussed in detail (Salvermoser et al., 2018). Additionally, other studies have shown the role of integrins in leukocyte migration; for example, a video-microscopy analysis in Peyer’s patches, showed a reduction in the recruitment and the adhesion of lymphocytes when they were treated with neutralizing antibodies against LFA-1 or the α4 subunit of integrins (Bargatze et al., 1995).

The reduction of recruitment of B cells to HEV was confirmed by homing assays, where it is observed that the loss of Myo1e causes a reduction of recruited B cells in the inguinal lymph node, concomitant with an accumulation of B cells in blood and spleen (Nolte et al., 2002, Ager, 2017). In our work we observed a reduction in the expression of integrins.

Of note, the role of Myo1e in the motility was only detected in activated B cells, since no significant differences were found in resting B cells. Activated B cells had reduced 2D migration, decreased adherence to fibronectin and hyaluronic acid; and diminished adherence to monolayers of bEnd.3 cells. Additionally, shorted membrane protrusions were found in Myo1e deficient B cells.

Migration is modulated by the expression of integrins; the cellular activation increased their expression (Chung et al., 2014) as well as chemokine receptors (Takabayashi et al., 2009, Goichberg et al., 2006), furthermore they are critical for the rearrangement of the cytoskeleton. Through different signaling pathways, these proteins contribute to the generation of membrane projections and as anchors to support the force needed for the motility. (Hood and Cheresh, 2002, Kritikou, 2008). Our results have shown that the loss of Myo1e in B lymphocytes causes a decrease in the expression of LFA-1.

The formation of “integrin cluster,” originates the autophosphorylation of FAK in Tyr^397^ (Calalb et al., 1996) and allows the recruitment of paxillin (Hu et al., 2014) tensin (Qian et al., 2009), and talin (Nader et al., 2016), which are necessary to form a complex that stabilizes the adhesion. According to our results, FAK associates with Myo1e in B cells; this interaction had been reported in WM858 melanoma cells (Heim et al., 2017) The interaction between FAK and Myo1e affects the autophosphorylation of FAK when CXCL12 stimulated the cells. Studies in DU-145 cells (human epithelial cells) and the hematopoietic precursors (HSC) demonstrated that CXCL12 modulates the phosphorylation of FAK and the expression of β3 and α5 integrins, which then contributes to the adhesion mediated by VCAM-1 (Engl et al., 2006, Glodek et al., 2007). Therefore, we hypothesize that Myo1e is responsible for carrying FAK towards the integrin, this allows FAK autophosphorylation causing the formation of a complex, which is necessary for efficient cell adhesion and motility.

The membrane protrusions are important morphological structures for motility and protein localization (Tanaka et al., 2017, Xue et al., 2010). The interaction of Myo1e with the CARMIL protein is critical for the formation and elongation of the actin filaments (Liang et al., 2009).. In this work, we demonstrated the association of CARMIL with Myo1e in B cells. This result suggests that Myo1e deficiency does not allow the recruitment of CARMIL causing the reduction of the membrane extensions.

The spreading is a mechanism used by the cell to maximize the contact area with different ligands. This is an essential step for slow rolling and then cellular transmigration. Deficiency in spreading has been described as a disturbance in the integrity of the cytoskeleton (Wakatsuki et al., 2003, Kim and Wirtz, 2013). In our work, we showed that Myo1e deficiency causes a reduction in spreading indicating that Myo1e is also involved in cell deformation.

RAC-1 is an essential small GTPase involved in the formation of actin filaments, spreading and cell motility. Its deficiency, in mouse embryonic fibroblasts (MEFs), has shown a reduction in the activity of RAC-1, altering cell morphology (Chang et al., 2011). This phenomenon has also been reported in the HeLa cell line, wherein the absence of RAP1, there is a decrease in the activity of Rac-1 affecting cell spreading and motility (Arthur et al., 2004)

The phosphorylation of AKT Thr^308^ regulates the activity of Rac-1 through PDK1 (Higuchi et al., 2008, Niba et al., 2013, Liu et al., 2018). There is an increase in the activity of RAC-1 and the phosphorylation of AKT by stimulation with growth factors, as in the cell line MDA-MB-231 (mammary gland epithelial cells) treated with epidermal growth factor (EGF) (Yang et al., 2011). Our results show a decrease in the spreading of activated Myo1e^−/−^ B lymphocytes that correlates with reduced activity of RAC-1 and reduced phosphorylation of AKT^Thr308^. FAK through its kinase activity allows the phosphorylation of PI3K. PI3K is an enzyme involved in the conversion of phosphatidylinositol 4,5, bisphosphate (PIP2) to phosphatidylinositol 3,4,5 triphosphate (PIP3) in the membrane (Agelaki et al., 2007, Matsuoka et al., 2012) PIP3 allows the anchorage of AKT to the membrane and promote the activity of RAC-1. The inhibition the activity of the FAK/PI3K/RAC-1 pathway, in the EA.hy926 cell line (human endothelial cells) and the MCF7 cell line (human epithelial cells) decreases cell migration (Kallergi et al., 2007, Huang et al., 2013). We found that in the absence of Myo1e, activated B cells stimulated with CXCL12 show a similar phenotype than wild type B cells inhibited by Ly294002 (a PI3K inhibitor). These results were corroborated with the reduction in the phosphorylation of FAK, AKT and the activity of RAC-1. As a whole, these results indicated that Myo1e is involved in the FAK/PI3K/RAC-1 signaling pathway. In conclusion, we have presented evidence that Myo1e is critical for the recruitment and adhesion of B cells to the inguinal lymph node through the localization of integrins and this phenomenon is modulated by the FAK/PI3K/RAC1 signaling pathway.

## Materials and methods

### Mice and reagents

We use female C57BL/6J or B6.129S6(Cg)-*Myo1e^tm1.1Flv^*/J (8–10 weeks; in all experiments). The mice were kindly provided by Dr. Richard Flavell (Yale School of Medicine, USA) and then bred and maintained in the animal facility at the “Centro de Investigación y de Estudios Avanzados” (Mexico City, Mexico) animal facility. The Animal Care and Use Committee of “Centro de Investigación y de Estudios Avanzados” approved all experiments.

All mice were allowed free access to water and a maintenance diet containing 20% protein (PicoLab^®^ mouse diet 20, LabDiet^®^ 5058, St. Louis, MO, USA) in a 12-hour light/dark cycle, with room temperature at 22 ± 2°C and humidity at 50 ± 10%. All cages contained Aspen chip and Aspen Shaving (50/50%) (NEPCO^®^ Warrensburg, NY, USA) as bedding, moreover, included wood shavings, bedding and a cardboard tube for environmental enrichment.

### Lymphocyte isolation and flow cytometry

Splenic mononuclear cells were isolated by Ficoll-paque Plus (GE Healthcare) (Little Chalfont, United Kingdom) density gradient separation, and then B220^+^ cells were enriched by panning, using plastic dishes coated with α–Thy-1 mAb ascites (NIM-R1) (Chayen and Parkhouse, 1982).

For activation, 2×10^6^ B cells were incubated in 1 ml 10% fetal bovine serum (FBS) (Thermo Fischer, Scientific) (Waltham, MA, USA) supplemented RPMI 1640 (Life Technologies) (Grand Island, NY, USA) containing LPS from Escherichia coli O55:B5 at 40 mg/ml (Sigma Chemical Co, St) (Louis, MO, USA) plus 10 U/ml IL-4 (R&D Systems) (Minneapolis, MN, USA) for 48 h at 37 □C and 5% CO2.

For immunostaining, we blocked the Fc receptors using 10% goat serum; the cell suspensions were immediately washed with PBS containing 1% BSA and 0.01% NaN3 (PBA). Depending on each experiment, one million cells were stained for 15 minutes using the antibodies described in the following section. After incubation, the cells were washed with PBA and were fixed with 1% Formaldehyde in PBS (0.5% Albumin, 0.01% Sodium azide, 100 ml PBS). The doublets were excluded with the gating on FSC-H vs. FSC-A, and the lymphocytes were identified by their scatter properties (FSC-A vs. SSC.A). The compensation was performed using single-stained cells for each of the fluorochromes used. The cells were evaluated using “BD LSRFortessa” flow cytometer (Becton-Dickinson) (San Jose, CA), and analyzed using FlowJo v.10 software (Tree Star, Inc.) (Ashland, Oregon). All the experiments were performed according to the flow cytometry guide-lines (Cossarizza et al., 2017).

### Antibodies and reagents

The antibodies used were: anti-B220-BV421 (clone RA3-6B2, BioLegend) (San Diego, California, USA), anti-B220/CD45R (clone RA3-6B2, BioLegend), anti-CD19 (Southern Biotechnology Associates) (Birmingham, Alabama, USA), anti-CD29 (clone hm *B*1-1, BioLegend), anti-LFA-1 (clone Hl111, BioLegend), anti-CD62L (clone DREG-56, BioLegend), anti-TLR-4 (clone TF901, BioLegend), anti-CXCR4 (clone 2G8, BioLegend), anti-CD44 (clone IM7, BioLegend), anti-Myo1e (clone PAD434 Cloud Corp) (Katy, TX, USA), anti-CARMIL (clone E-10, Santa Cruz, Biotechnology) (Dallas, TX, USA), anti-WASp (Clone EP2541, Abcam) (Cambridge, UK) anti-RAC-1 (clone B-8, Santa Cruz, Biotechnology), anti-RAC-1 GTP (clone 26903, ser-61, Biomol) (Hamburg, Germany), anti-PI3K (clone sc-1637, Santa Cruz, Biotechnology) anti-AKT (clone sc-5298, Santa Cruz, Biotechnology), anti-phospho-AKT (clone sc-271966, Santa Cruz, Biotechnology). Other reagents included, TRITC-Phalloidin (Thermo Fischer, Scientific), Hoescht 33342 (Thermo Fischer, Scientific), Ly294002 (Sigma Aldrich), The murine CXCL12 was purchased from PeproTech (Rocky Hill, NJ, USA).

### Immunofluorescence microscopy

Cells were fixed 20 minutes with paraformaldehyde at 4%. After washing, the cells were permeabilizated 30 minutes with Triton X-100 (0.1%). Then, the Fc receptors were blocked with goat serum to avoid nonspecific binding. Immunolabeling with primary antibodies was performed by 30 minutes incubation at 4 C, followed by washing and incubation with species-specific fluorescence-labeled secondary antibodies or TRITC-phalloidin (Thermo Fischer, Scientific). The preparations were mounted with Vecta-shield (Cat. H-1000 Vector labs) (Burlingame, CA, USA). The slides were analyzed in confocal microscopy (Leica Microscopy, TCS SPE, Model DMI4000) (Wetzlar, Germany). Quantification of intensity fluorescence was performed using the program LAS AS lite 5.0 (Leica Microscopy).

### Homing assays

B cells purified from spleen from Myo1e^+/+^ or Myo1e^−/−^ mice were labeled with 0.1 μm or 0.6 μm of Carboxyfluorescein succinimidyl ester (CFSE) (Thermo Fischer, Scientific), respectively, or vice versa, in a complementary set of experiments. The cells were mixed at different ratios: 25%, 50% or 75% Myo1e^+/+^ B cells with the respective percentage of Myo1e^−/−^ B cells to complete 100%. The mixed suspensions of 1 x10^7^ B cells were injected via the tail vein. One-hour previously, the right inguinal lymph node of a host wild type mice was inoculated with CXCL12 (25 ng/ml), while the left inguinal lymph node was inoculated with PBS. The host mice were sacrificed 2 hours after inoculation. The blood, the spleen, and the inguinal lymph nodes were extracted. After that, the cells were recovered and measured using “BD LSRFortessa” flow cytometer (Becton-Dickinson) (San Jose, CA, USA), and analyzed using FlowJo v10 software (Tree Star, Inc.) (Ashland, Oregon, USA). For intravital microscopy, both inguinal lymph nodes were extracted to quantify the numbers of cells.

### In vitro chemotaxis assays

For quantification of migration, a Zigmond chamber (Neuroprobe) (Gaithersburg, MD, USA) was used. Briefly, one million of activated B lymphocytes from Myo1e^+/+^ and Myo1e^−/−^ mice, were suspended in 0.5 mL of 10% FBS supplemented RPMI 1640 (Life Technologies) and immediately plated onto glass coverslips, previously coated with fibronectin (2.5 μg/mL) (Sigma-Aldrich), that were incubated for 30 min at 37°C and 5% CO2, to allow the cells to attach. The coverslips, with the cells attached, were gently washed with PBS, One of the grooves in the Zigmond chamber was filled with supplemented medium (approximately 70 μL), and the other groove was then filled with CXCL12 (2.5 μg/uL) (PrepoTech) also dissolved in a supplemented medium. A baseline image was obtained at 10× magnification, and digital images of the cells were taken every 30s for 1h maintaining the temperature of the room between 35 and 39°C. For analyzing the trajectories and speed of migration, the migration tracks were traced for at least 100 lymphocytes of Myo1e^+/+^ and Myo1e^−/−^, in five independent experiments, using the NIH ImageJ software with chemotaxis and migration tool 2.0 (Ibidi, Martinsried, Munich, Germany) (Gorelik and Gautreau, 2014).

### Adhesion assays

Polystyrene plates with 96 wells (Nalge Nunc International) (Penfield, NY, USA) were coated with Hyaluronic acid (2.5 mg/ml) (Sigma-Aldrich), fibronectin (2.5 mg/ml) (Sigma-Aldrich) or poly-lysine (0.01%) (Sigma-Aldrich), 1 hour at 37 C. After incubation, the plates were washed twice with PBS before adding 4 × 10^5^ panning-enriched B cells in 200 μl of RPMI 1640 per well. The cells adhered for one h at 37°C and then, the plates were washed with PBS. The cells were fixed 10 min with 4% paraformaldehyde, before adding crystal-violet (7.5 g/l crystal-violet, 2.5 g/l NaCl, 1.57% formaldehyde, 50% methanol) for an additional 5 minutes. After that, the plates were solubilized with 10% SDS, and the amount the remaining dye in the plates was registered at 540 nm (Multiskan Ascent) (Thermo Fischer Scientific). Non-specific dye bound to empty wells was subtracted, and the absolute binding was calculated. The absorbance was determined in four wells per condition.

### Western Blot

B cells were lysed with RIPA buffer (20 mM Tris–HCl (pH 7.5), 150 mM NaCl, 1 mM, EDTA, 1 mM EGTA, 1% Triton X-100, 1 μg/ml leupeptin, 10 μg/ml aprotinin, and 1 mM PMSF) 30 min at 4°C. The protein content was determinate with the Modified Lowry Protein Assay Kit (Thermo Fischer, Scientific). Proteins were separated via 12% SDS-PAGE at 85 V, and 50 μg of protein was added from each sample to independent wells in the gel. After electrophoresis separation, the proteins were transferred to a nitrocellulose membrane (BIO-RAD) (Hercules, CA, USA) at 120 V, 1.5 hours. After transference, the membranes were blocked 30 minutes with albumin serum bovine (BSA) (5%) (Thermo Fischer, Scientific) After blocking; the membranes were incubated one hour at 37 °C with specific antibodies. After washing with TBS-Tween 20 (0.01%) (Sigma Aldrich), the membranes were incubated with the respective secondary HRP-labeled antibody. Finally, the blots were revelated with Western Blotting Chemiluminescence Luminol Reagent (Santa Cruz, Biotechnology). Tubulin or actin was used as loading controls.

### Intravital Microscopy

Myo1e^+/+^ host mouse was anesthetized by intraperitoneal injection of 12.5 mg/kg xylazine and 125 mg/kg ketamine hydrochloride (Sanofi, Mexico-City, Mexico). Then, the inguinal lymph node of the was inoculated with CXCL12 (25 ng/ml) (PeproTech). One hour later, 1×10^7^ Hoestch 333462 labeled B cells were directly injected via the carotid artery. The venules of the inguinal lymph node were recorded using the an intravital upright microscope (Axioscope, Model A1, Zeiss), (Oberkochen, Baden-Württemberg, Germany) with a 40 x and 0.75 saline immersion objective (Zeiss, Microscopy) (Oberkochen, Baden-Württemberg, Germany) Videos and images were analyzed using ImageJ (NIH, Bethesda, MD. USA) and Zen Blue Edition 2.5 software (Zeiss, Microscopy). The diameter of the venules, the number of adherent cells, the number of transmigrated cells and the velocity of the cells were measured with ImageJ. The cells flux, the blood flux, the slow rolling and the rolling were analyzed by Zen Blue Edition 2.5 software (Zeiss, Microscopy).

### Pharmacological inhibition treatment

Ten million activated B cells treated (2 h) with 20 μM LY294002 (Sigma Aldrich), were stimulated with CXCL12 (PeproTech) in RMPI-1640, supplemented with 5% fetal bovine serum or only supplemented medium. After blockage and stimulation, the cells were used in different experiments as indicated.

### Co-Immunoprecipitation assay

Protein extracts (500 μg) of resting or activated B cells were used, the lysates were centrifuged 18,000 g, 30 min at 4°C. The supernatants were mixed with anti-Myo1e, anti-Focal adhesion kinase (FAK) or anti-CARMIL, using rabbit IgG or rat IgG, as isotype controls, respectively. The supernatants were incubated overnight at 4°C in agitation; then, the complexes were precipitated with protein G-agarose (Life Technologies), maintaining the temperature at 4°C. Complexes were washed three times with RIPA buffer and boiled in Laemmli buffer. SDS-PAGE and western blotting was performed as previously indicated.

### Statistical analysis

Data are presented as the arithmetic mean with standard deviations; the *t* Student test was used for evaluating statistical differences. A p-value of <0.05 was considered statically significant. The p values are represented as *p<0.05, ** p<0.01 and ***p<0.001 and the number of samples or cells (n) used are mentioned in each figure legend.

## Acknowledgments

D.AG-P. designed and performed the experiments; and wrote the paper. E-V designed and performed intravital experiments, M-S designed the experiments, supervised the work, LS-A designed the experiments, supervised the work, and wrote the paper.

The authors thank Dr. Richard Flavell, for the kind donation of the Myo1e ^−/−^ mice; Dr. Santiago Partida-Sanchez for his help to reinitiate our colony of mice; Dr. Hector Romero-Ramirez, Mr. Lenin Estudillo, and M.Sc. Itze Cecilia Navarro Hernandez for their support; and, MVZ Ricardo Gaxiola-Centeno for taking care of the mice

